# Not just with the natives, but phylogenetic relationship between stages of the invasion process determines invasion success of alien plant species

**DOI:** 10.1101/2022.10.12.512006

**Authors:** Achyut Kumar Banerjee, Hui Feng, Xinru Liang, Fengxiao Tan, Jiakai Wang, Yuting Lin, Minghui Yin, Hao Peng, Wuxia Guo, Nannan Zhang, Yelin Huang

## Abstract

Darwin proposed two alternate hypotheses on the invasion success of alien plant species based on their close or distant phylogenetic relationship with the natives. Here we tested these hypotheses along the invasion process at large spatio-temporal scale. Using two abundance-based phylogenetic metrics for the native and alien flora of China, our study showed that invasion success of alien plant species is influenced by the phylogenetic relationship with native and alien species of different invasion stages. Phylogenetic similarity with the native species helps the alien ones to introduce and naturalize, but phylogenetic dissimilarity with the natives facilitates invasion success. The co-occurrence of invasive and naturalized aliens also formed more clustered assemblages, showed specialized responses to environmental stress, and provided temporal stability to the phylogenetic measures. The native-alien phylogenetic relationship is dynamic across spatial, environmental, and taxonomic scales. Therefore, assessing Darwin’s naturalization conundrum at different gradients of community assembly process is important.

## Main Text

Biological invasions are considered one of the most important global problems experienced by ecosystems^1^. Phylogenetic relatedness of the alien species to the existing community members is hypothesized to either facilitate (pre-adaptation hypothesis^2^) or hinder (Darwin’s naturalization hypothesis ^3^) the successful establishment of alien species in the community. It has been argued that these two seemingly contradictory hypotheses, also known as Darwin’s naturalization conundrum^4^, can address the community assembly process at different spatial scales. Numerous studies have investigated the phylogenetic relationship between native and alien species to provide evidence supporting these two hypotheses^5^.

At a large spatial scale, phylogenetic relationship as an invasion correlate has largely been examined for alien species of different invasion stages together ^6^ rather than considering them separately in a single analysis. Biological invasion is a process that can be divided into a series of stages referring to an alien species depending on where in the invasion process it is – introduced, naturalized, and invasive ^7^. Few studies that have examined the phylogenetic relationship across the invasion continuum ^8^ have provided clues that Darwin’s naturalization conundrum can be addressed by considering the relationship between native and alien species of different invasion stages. By considering cultivated alien plant species of South Africa, Omer et al. ^9^ revealed that the probability of introduction increased with higher phylogenetic distance from the natives (naturalization hypothesis), introduced species phylogenetically similar to the native species could successfully naturalize (pre-adaptation hypothesis), and successful invaders need to be functionally different than the native species (naturalization hypothesis). By comparing functional traits based on phylogenetic history between the native and alien species in temperate Central Europe, Divíšek et al. ^10^ also reported a similar phylogenetic pattern.

However, large-scale phylogenetic patterns result from the interaction between ecological and evolutionary processes functioning at different spatial and temporal scales ^11^. Therefore, the inferences of previous studies are often restricted either by considering different invasion stages together ^6^ or by the spatial extent of the study area with limited environmental heterogeneity ^9^. Moreover, interactions between alien species can also influence the community assembly process ^12^, and phylogenetic evidence of such interactions is rare. We were interested in exploring the phylogenetic relationship of alien plant species not only with natives but also between different stages of the invasion process and how far the previous observations on Darwin’s naturalization conundrum can remain consistent at large spatio-temporal scales. We conceptualized four objectives to test these objectives across the invasion continuum along environmental, temporal, and taxonomic scales.

### Hypothesis 1

In line with previous observations of Omer et al. ^9^ and Divíšek et al. ^10^, we first hypothesized that the phylogenetic relationship between native and alien species would vary depending on the invasion stages.

### Hypothesis 2

In response to environmental stress, phylogenetically closely related species should form a more clustered assemblage, as ecological traits are often phylogenetically conserved, and very few clades have evolved to survive in harsher environmental conditions ^13^. The phylogenetic clustering should vary across the invasion continuum as the alien species of different invasion stages can respond differently to the environmental filtering process ^14^.

### Hypothesis 3

Since phylogenetic composition reflects adaptations shared in evolutionary history ^15^, the phylogenetic change pattern will be influenced by the deterministic processes (i.e., biotic interactions) ^16^. We expected that the communities with the introduced species show relatively stable phylogenetic changes (due to phylogenetic similarity with the natives, pre-adaptation hypothesis), and the invasive and naturalized aliens would create more phylogenetic changes (due to phylogenetic dissimilarity with the natives, naturalization hypothesis) in the community over time.

### Hypothesis 4

The phylogenetic pattern can vary depending on the taxonomic scope (whole flora or family-specific) of the study ^6^. The phylogenetic pattern should be more pronounced at smaller extents, where species of the same family are more likely to respond to environmental gradients similarly. Because individual families have different evolutionary histories, the phylogenetic patterns would vary among them.

We tested these hypotheses by focusing on the native and alien plant species of China, one of the 12 megadiverse countries in the world with a large geographic extent and wide environmental variations. The history of alien plant introduction into China goes back more than 2000 years, and many of them have become naturalized and subsequently invasive ^17^. With increased globalization and the gradual opening of China’s economy to the global markets in the 21^st^ century, the novel introduction of alien species is expected to increase. Therefore, understanding the phylogenetic relationships between the alien species of different stages of the invasion continuum and their influence on the community assembly process have crucial management implications.

A recent study has assessed the phylogenetic relationship between native and alien plant species of China at a large spatial scale ^18^; however, the pattern and its dynamics along the invasion continuum are still to be explored. Our objectives of this study were two folds: 1) to assess the phylogenetic relationship between native and alien species of different invasion stages (introduced, naturalized, and invasive), and 2) to check whether the phylogenetic pattern will change across environmental, temporal, and taxonomic scales. We addressed these objectives using a highly resolved phylogenetic tree and abundance-weighted phylogenetic measures between native and alien plant species of China.

## Results

A total of 8,65,752 occurrences for 19,674 species (Fig.1a, Supplementary Data 1) was used to estimate the two phylogenetic measures, net relatedness index (NRI) and the nearest taxon index (NTI), across 57 ecoregions and seven periodic intervals based on a highly resolved phylogenetic tree (Extended Data figure 1). The number of ecoregions with co-occurring native and alien species (see Fig.1b for the abbreviations) increased over periodic intervals (Supplementary Table 1).

**Fig. 1:**
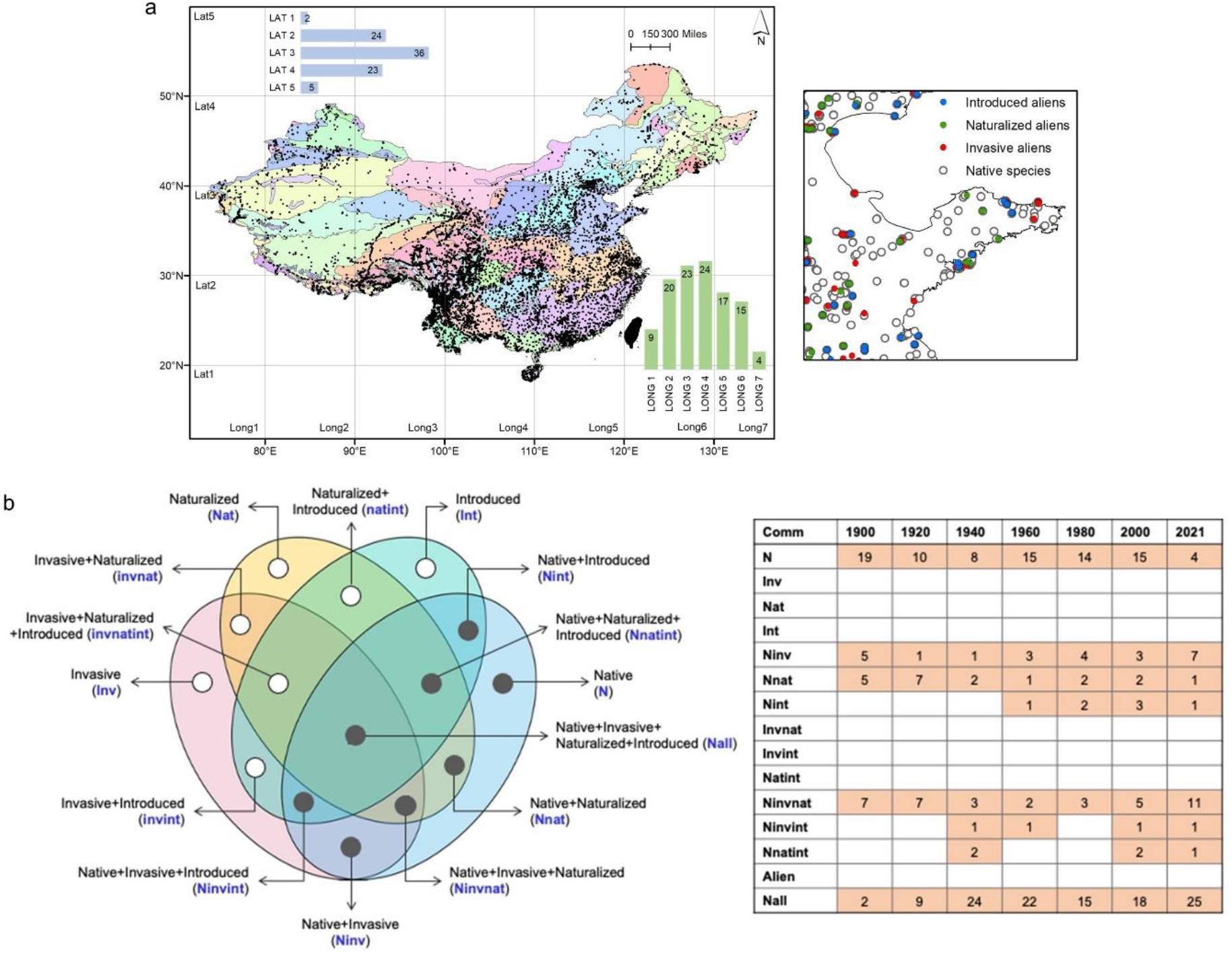
a) Map of the study area showing the distribution of native and three categories of alien species (introduced, naturalized, and invasive) across 57 ecoregions present in China. The bar plots show the number of ecoregions in each latitudinal and longitudinal grid at 10-degree intervals. b) The Venn diagram shows 15 possible community compositions (and their abbreviations) having the native species and three categories of alien species, and the number of species at each of the seven periodic intervals.

### Phylogenetic patterns

At the species level, the average phylogenetic measures (averaged for the eight communities and seven periods) were positive (Table 1). The NRI values were mostly higher than the NTI values, except for the Ninvint, Ninvnat, and Nnat communities. The average phylogenetic measures at the family level were significantly lower (at p = 0.05 level) than that observed at the species level, except for the NRI values for the Poaceae (Table 1; Supplementary Table 2).

**Table 1:**
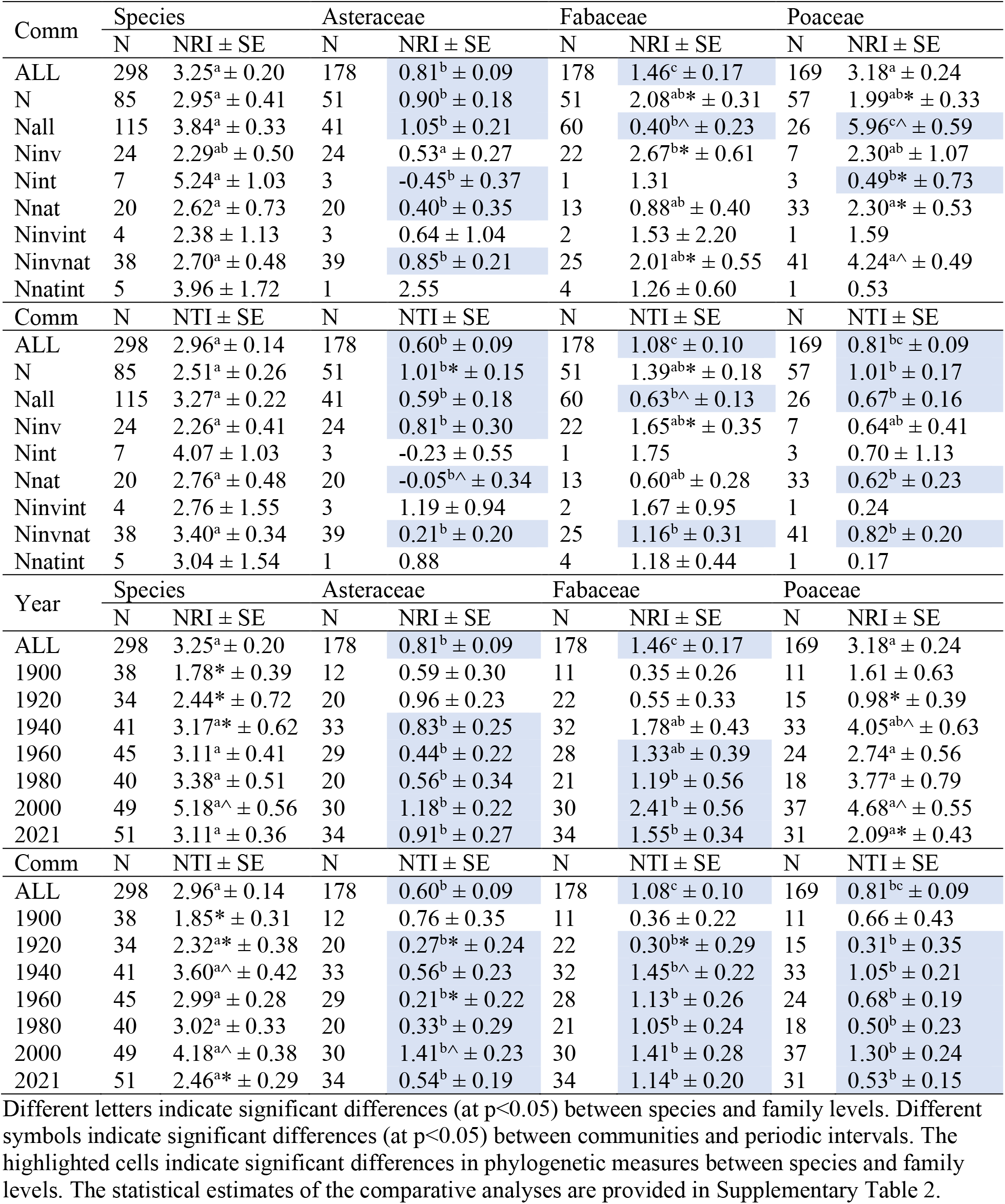
Average net related index (NRI) and nearest taxon distance (NTI) values for eight communities (Comm) and seven periodic intervals (Year) at species and family levels. The communities and periodic intervals are identified in text and Fig. 1.

Considering the communities (averaged for the seven periods), the Nint and Ninv communities had the highest and lowest phylogenetic measures, respectively. The average NRI and NTI values of the Nnatint and Ninvnat communities were higher than the communities where the naturalized and invasive aliens occurred only with the native species (Table 1). Among the three families, the maximum number of significant differences from species level was observed for Asteraceae, and marginally higher NRI values were observed for the Ninv communities of Fabaceae and Poaceae (Table 1). Significant differences were also observed between the phylogenetic measures of the eight communities. The correlation analysis revealed very few significant correlations between the phylogenetic measures of the eight communities, both at species and family levels (Supplementary Table 3).

When the periods were considered (averaged for the eight communities), significant differences were observed between the phylogenetic measures of the seven periodic intervals (Supplementary Table 2). The posthoc tests showed significantly higher NRI and NTI values at both species and family levels in the 1981-2000 periodic interval (Table 1). The phylogenetic measures of all three families were significantly lower than that observed at the species level since 1920 (for NTI), except for the NRI values for the Poaceae.

### Phylogenetic anomalies

At the species level, the phylogenetic anomalies (PAμ, averaged for the seven communities) were positive (PA_NRI_: 0.014±0.24; PA_NTI_: 0.53±0.23) and significantly different from the negative PA_NTI_ values observed at the family level (Supplementary Table 4). The alien species of three invasion categories generated different PAμ patterns (Fig.2a). The average PAμ values were positive for the Nint communities. In the presence of naturalized aliens in the community (Nnat), the average PA_NRI_ anomaly was negative, and the PA_NTI_ anomaly was positive. The invasive aliens occurring alone (Ninv) created negative PAμ values. However, positive PA_NRI_ and PA_NTI_ were observed in the Ninvnat community, and positive PA_NTI_ was observed in the Ninvint community. The PA_NTI_ of the Ninvnat community was significantly higher than the null value at the species level (Fig.2a). Significant differences in the PAμ values between the communities were observed at the family level (Fig.2b-d).

**Fig. 2:**
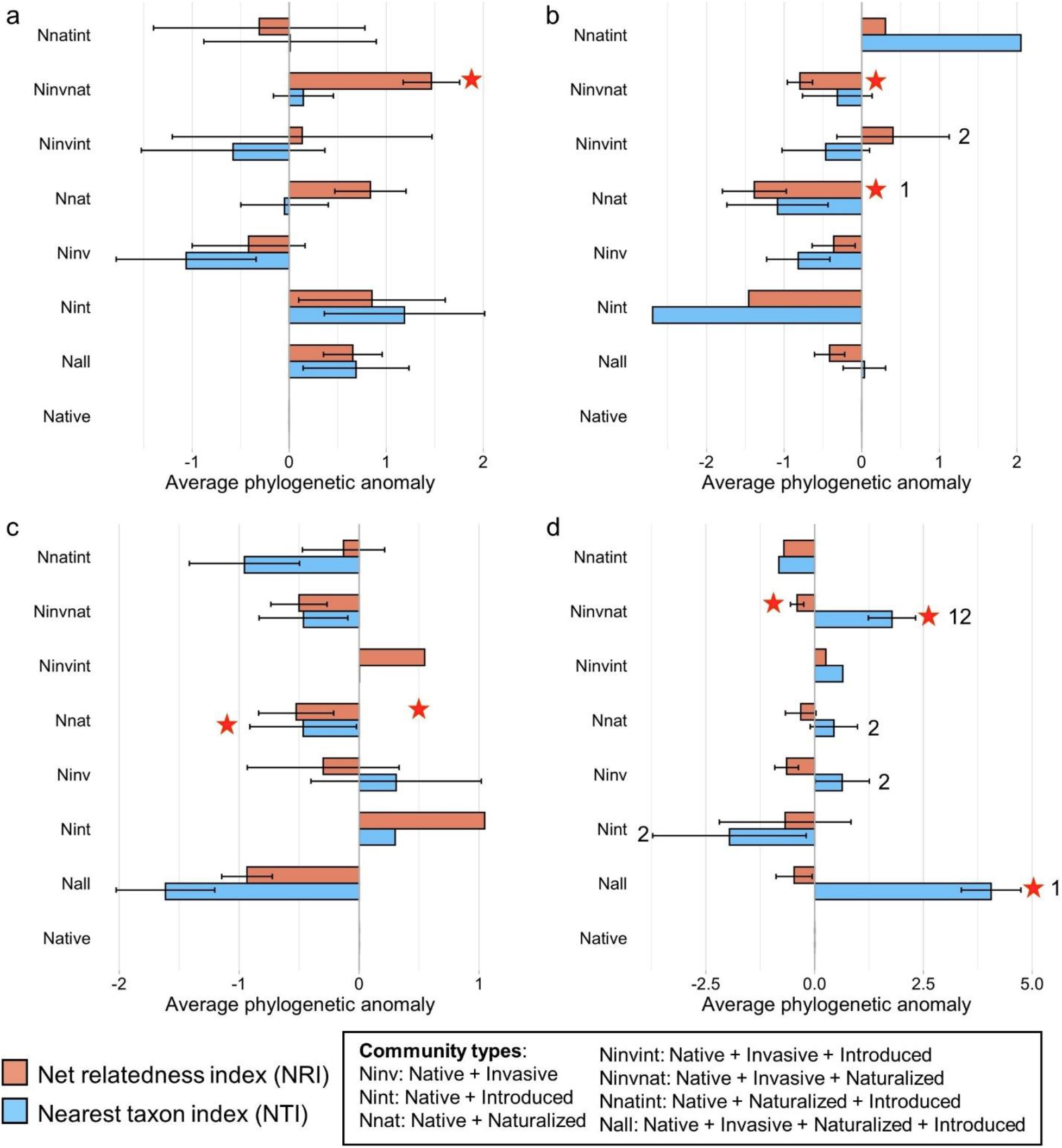
Phylogenetic anomaly of the two measures (NRI - net relatedness index and NTI - nearest taxon index) caused by the introduction of alien species a) at the species level, and for b) Asteraceae, c) Fabaceae and d) Poaceae. The bars represent mean phylogenetic anomaly values with standard errors. The red stars indicate a significant difference (at p<0.05 level) in the phylogenetic anomaly values from zero (i.e., communities with native species only). Different letters above the bars indicate significant differences in phylogenetic anomaly values between the communities. The statistical estimates of the comparative analyses are provided in Supplementary Table 4.

### Phylogenetic changes over time

The phylogenetic change over time (PTμ, averaged for the eight communities) was more for the NRI values (PT_NRI_: 0.21±0.2) compared to the NTI values (PT_NTI_: −0.05±0.2). Among the eight communities, higher PTμ values were observed for the communities having the introduced aliens (Nint, Ninvint, Nnatint) (Fig.3a). At the family level, the highest positive change was observed for the family Fabaceae, especially for the communities with the invasive aliens (Ninv, Ninvnat) (Fig.3c). Presence of invasive aliens of Asteraceae, on the other hand, created negative phylogenetic changes over time (Fig.3b). However, we found no significant differences in the PSμ values between the communities, both at species and family levels (Supplementary Table 5).

**Fig. 3:**
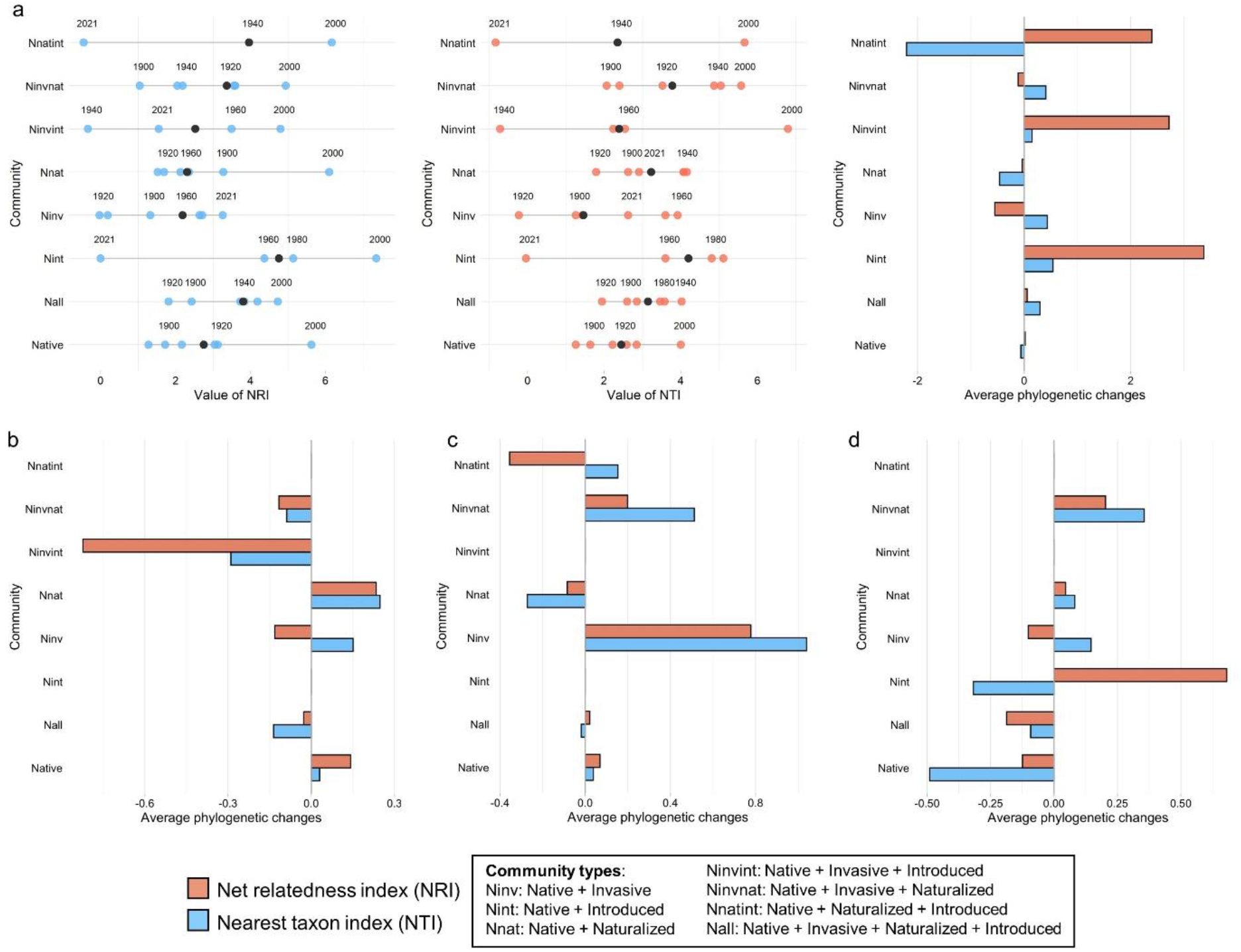
Changes of the phylogenetic measures (NRI - net relatedness index and NTI - nearest taxon index) over time – a) the beam plots showing the NRI and NTI values at each of the seven periodic intervals with the median values marked in black for each of the eight communities; the bar plots represent average phylogenetic changes over time at the species level, and for c) Asteraceae, d) Fabaceae, and e) Poaceae.

### Phylogenetic changes across space and environment

The average phylogenetic measures (averaged for the communities) were significantly different along the latitudinal and longitudinal gradients, both at species and family levels (Supplementary Table 6). The posthoc tests showed that at the species level, the average phylogenetic measures were significantly higher in the grids south of 40^0^N and west to 110^0^E along the latitudinal and longitudinal gradients, respectively (Fig.4). However, for Fabaceae and Poaceae, the latitudinal and longitudinal thresholds shifted to 30^0^N and 100^0^E, respectively. Significantly higher NRI and NTI values were observed in grids north of 30^0^N for Fabaceae and south of 30^0^N for Poaceae. The phylogenetic measures of the native species communities were more influenced by the latitudinal and longitudinal gradients (Supplementary Table 6), and more significant differences were observed along the longitudinal gradient compared to the latitudinal gradient. The logistic regression models showed that the phylogenetic measures were negatively influenced by annual mean temperature (BIO1) across the eight communities (Fig.5; Supplementary Table 7). More significant relationships between phylogenetic measures and bioclimatic variables were observed at the family level, especially for the Ninvnat communities of the three families (Fig.5). In general, NRI and NTI values were negatively influenced by BIO1 and positively by annual precipitation (BIO12) across the communities.

**Fig. 4:**
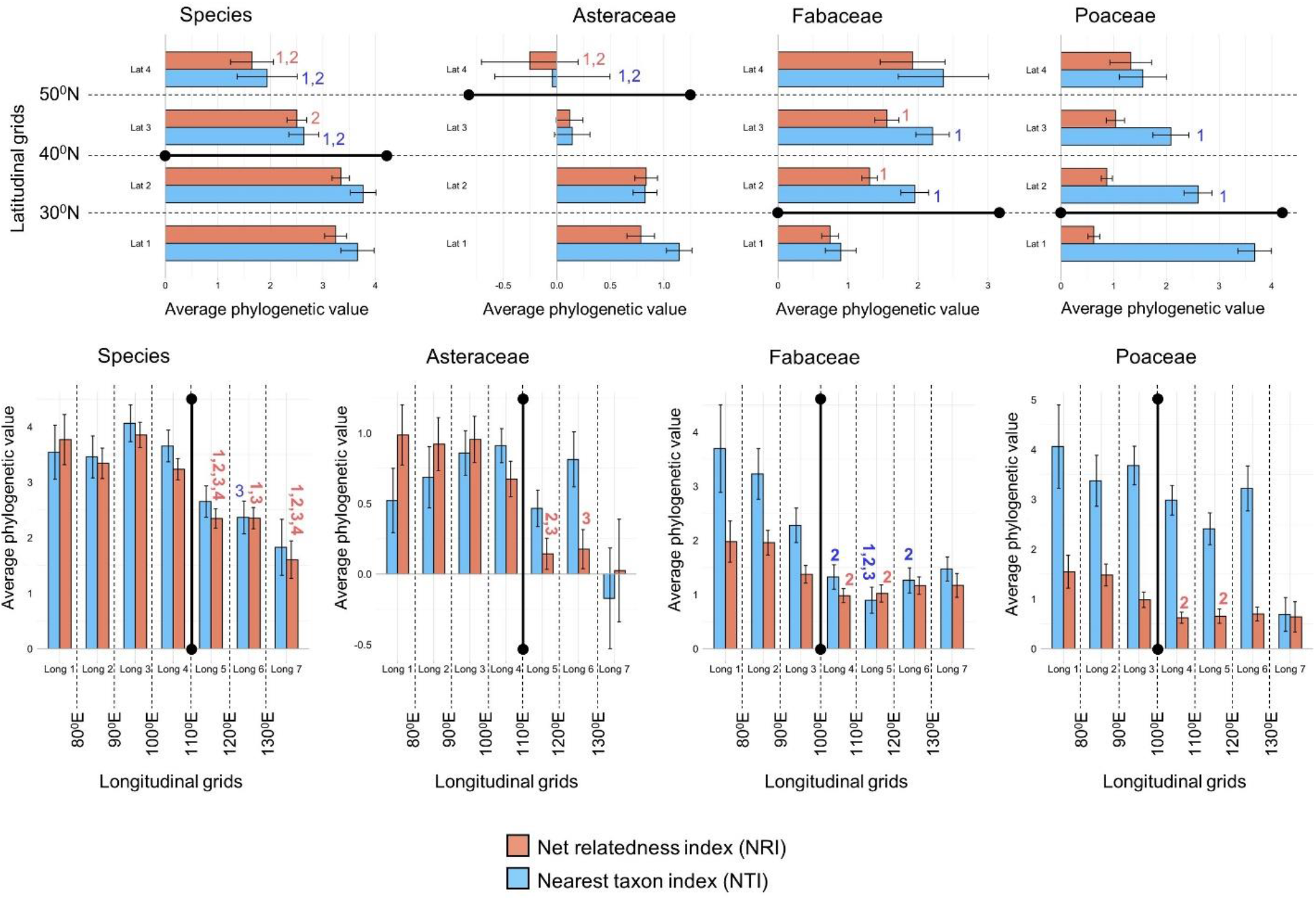
Variations of the two phylogenetic measures (NRI - net relatedness index and NTI - nearest taxon index) across latitudinal (top) and longitudinal (bottom) gradients. The bars represent mean NRI and NTI values with standard errors. Different numbers above the bars indicate significant differences between the grids separated by 10-degree intervals. The grids are identified on the study area map in Fig. 1. The statistical estimates of the comparative analyses are provided in Supplementary Table 6.

**Fig. 5:**
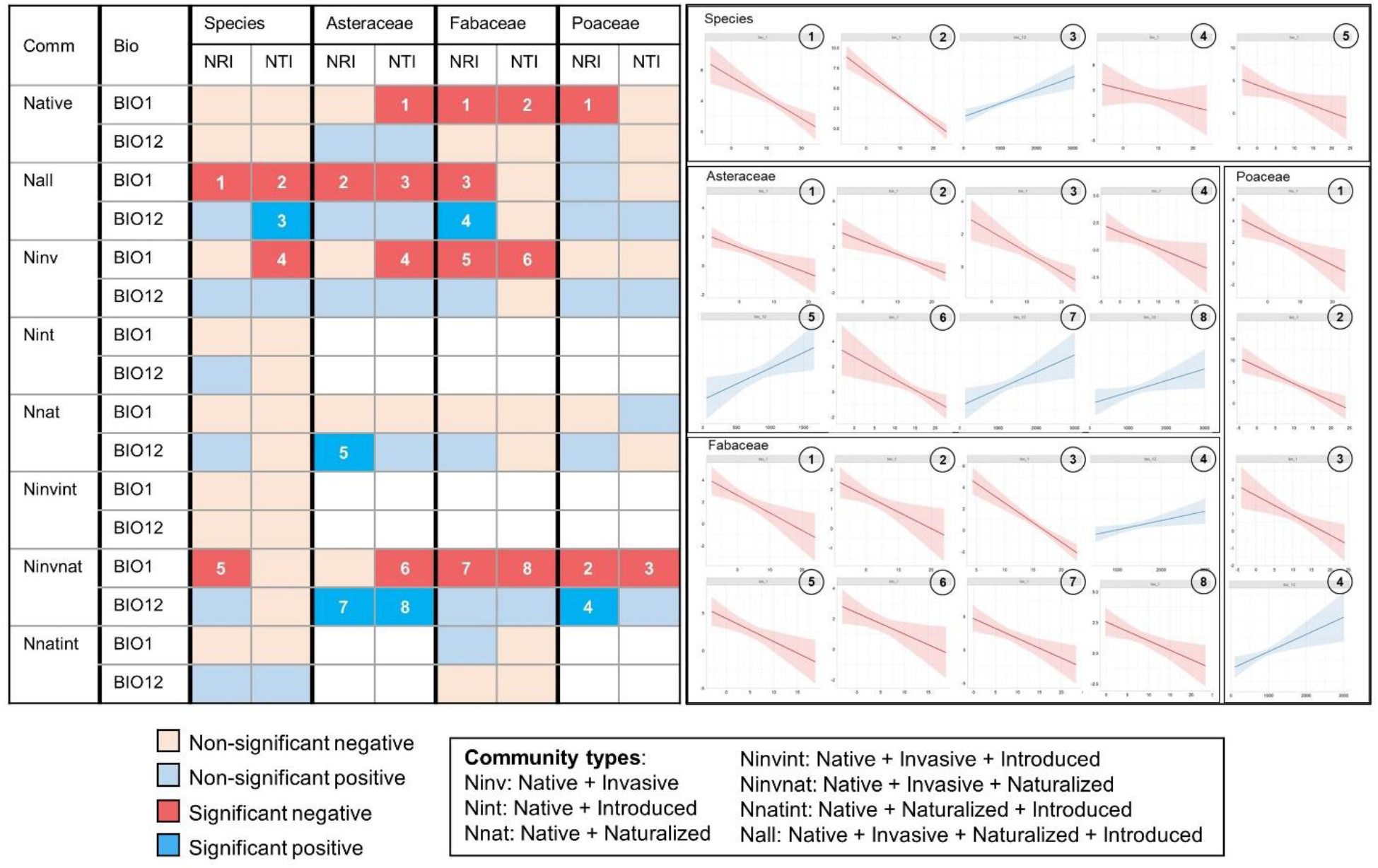
Matrix of regression analysis outputs showing the influence of the two bioclimatic variables (BIO1 and BIO12) on the net relatedness index (NRI) and nearest taxon index (NTI) of the eight communities at species and family levels. The regression plots show the significant relationships (numbers corresponding to the matrix) between the bioclimatic variables and predicted values of the phylogenetic measures with 95% confidence intervals. The statistical estimates of the regression models are provided in Supplementary Table 7.

## Discussion

Our study found positive phylogenetic measures for all seven alien communities, indicating that the existing lineages of the alien species may lead to small mean pairwise distances in the community (positive NRI), and the alien species often have a close relative in the community (positive NTI). These findings resonate with the pre-adaptation hypothesis, which posits that the phylogenetic relatedness of the alien species to the existing community members can facilitate their successful invasion ^2^.

However, the pattern of phylogenetic relationships varies across the communities depending on the position of the alien species in the invasion process, thereby supporting our first hypothesis. The communities having the introduced (Nint) and invasive (Ninv) aliens co-occurring only with the natives represented two edges of the phylogenetic spectrum. Firstly, the highest NRI and NTI values of the Nint communities indicate that for successful introduction into a native community, alien species should have an existing lineage or a close relative present in the community. This pattern supports the pre-adaptation hypothesis and contradicts the findings of Omer et al. ^9^. Secondly, in the communities with naturalized species, we found NTI>NRI in Nnat and NRI>NTI in Nnatint communities. This phylogenetic pattern indicates that for successful naturalization, the alien species needs to have a close relative among the natives or existing lineages among the introduced species. This finding supports the pre-adaptation hypothesis and concurs with the findings of both Omer et al. ^9^ and Divíšek et al. Finally, the invasive aliens showed the maximum phylogenetic distance from the natives, thereby supporting the naturalization hypothesis ^3^. However, among the communities with the invasive species, the highest NRI and NTI values were observed in Ninvnat (native + invasive + naturalized) community, suggesting that the probability of successful invasion of an alien species increases in communities having naturalized species of the same genetic lineage or close relative of that.

The pattern of phylogenetic anomalies also varied across the invasion continuum. The presence of introduced species created a more clustered community (positive anomalies for both NRI and NTI). The naturalized species created clustered assemblage with their phylogenetically close relatives (positive anomalies for NTI). These findings match with previous studies, which reported increased phylogenetic clustering with the introduction of alien species in China ^18^ and North America ^19^. However, the invasive aliens caused community disassembly (negative anomalies for NRI and NTI) when they occurred alone with the natives and formed clustered assemblage when they co-occurred with the naturalized and introduced aliens. Furthermore, higher community assembly in Ninvnat communities than that of Ninvint suggested that the invasives are more phylogenetically related to the naturalized aliens than the introduced alien species. We also observed a significant correlation between the NTI values of the invasive and naturalized alien species communities. These findings show that the phylogenetic relationship between native and alien species changes from one invasion stage to another.

The native-alien phylogenetic relationship across the invasion continuum was also influenced by the environmental filtering process ^20^, as evident from the significant relationships between the environmental factors and phylogenetic measures only in the presence of the alien species in the community. The negative relationships between phylogenetic measures and annual mean temperature further indicated that with increased environmental stress, phylogenetic clustering increased between species sharing similar environmental space ^21^, as also observed for other plant ^22^ and animal ^23^ species, thereby supporting our second hypothesis. Along the invasion continuum, significant relationships between temperature and NTI were found in the communities with the invasive aliens and between temperature and NRI in Ninvnat communities. This phylogenetic pattern implies a more specialized response of the invasive and naturalized aliens to the environmental filters, possibly due to their ability to retain niche-related traits through evolutionary history. The finding thus provided phylogenetic support to the niche conservatism hypothesis ^24^, which has been reported previously for both plant ^25^ and animal ^26^ invasive species.

The effect of environmental filtering on the phylogenetic relationship was further influenced by spatial gradients of community assembly. Our study revealed that above 40^0^N, both the NRI and NTI values decreased significantly. The ecoregions above this latitudinal threshold are comprised of deserts (e.g., Taklamakan Desert), semi-desert areas (e.g., Alashan Plateau Semi-Desert), and dry grasslands (e.g., Mongolian-Manchurian Grassland). The environmental unsuitability of these ecoregions to sustain diverse life forms can reduce the chances of finding similar genetic lineages and closely related species ^27^, leading to reduced phylogenetic measures. In addition, a significant reduction of NTIs above 110^0^E indicates compositional changes of closely related species along the longitudinal gradient, similar to the pattern observed in the Eurasian forest-steppe region ^28^. Overall, the phylogenetic pattern along the latitudinal and longitudinal thresholds closely matched the biogeographic boundaries in China identified in previous studies based on floristic composition ^29^.

The native-alien phylogenetic relationship also varied across the temporal scale. The native communities were found to have the most stable phylogenetic changes over time, which increased with the presence of introduced aliens in the community. This finding refutes our third hypothesis and suggests that the close relationship between native and introduced species (i.e., the pre-adaptation hypothesis) is not stable at a temporal scale. The number of novel introductions has increased in China over time, especially since 1980, due to the country’s reformation and economic activities ^30^. The gradual opening of the country to international trading might have aided the introduction and spread of many alien species ^31^, leading to changes in the native-alien phylogenetic relationships. Unlike the introduced aliens, the invasive and naturalized species have lag periods to establish in the community ^32^ and, therefore, showed comparatively stable phylogenetic changes over time. Interestingly, among the seven alien species communities, the co-occurrence of the naturalized and invasive aliens in the community (Ninvnat) provided maximum temporal stability to the phylogenetic measures. This pattern proves the pre-adaptation hypothesis in communities with invasive and naturalized aliens.

Deep down the tree, at the family level, the phylogenetic pattern varied from that observed at the species level and between three families, thereby supporting our fourth hypothesis. Firstly, we observed less clustered assemblages at the family level, thereby contradicting the previous studies, which considered the three alien species categories together to decipher the native-alien phylogenetic relationship ^6^. Secondly, the order of communities based on phylogenetic relatedness between native and alien species varied from that observed at the species level (i.e., the Nint and Ninv communities representing two ends of the high-low spectrum). Therefore, pre-adaptation and naturalization hypotheses explaining the influence of phylogenetic relationships on the invasion continuum varied at the family level. For example, negative phylogenetic measures were observed only in the Nint communities of Asteraceae, indicating that the presence of phylogenetically close native species or native species of similar genetic lineages in the community may hinder the successful establishment of Asteraceae aliens. This phylogenetic pattern, therefore, supports Darwin’s naturalization hypothesis ^3^. Further, we observed the highest NRI in the Fabaceae Ninv community, which indicates that the presence of native species of a similar genetic lineage may influence the successful invasion of a Fabaceae alien species, thereby supporting the pre-adaptation hypothesis. The pattern was also evident from the phylogenetic anomaly values, as the introduction of Fabaceae invasives was found to create more clustered assemblages ^33^. Thirdly, more phylogenetic changes over time were observed in communities with the invasive aliens at the temporal scale. From the evolutionary perspective, the phylogenetic structure is heavily influenced by polyploidy, which occurs more often in Poaceae than in any other angiosperm family ^34^, which might explain the high amount of phylogenetic changes over time, even among the natives of this family. Fourthly, with the decrease in the phylogenetic resolution, we observed more significant relationships between the phylogenetic measures and bioclimatic variables, especially in Asteraceae and Fabaceae communities. The pattern is expected since the species of the same family are expected to respond similarly to the environmental gradients ^35^. In addition, the spatial patterns of phylogenetic variation also changed for Fabaceae and Poaceae communities with the latitudinal and longitudinal thresholds shift. These findings provide evidence of the varied environmental influence at different levels of the phylogenetic tree, often obscured by increased phylogenetic extent by combining species with different response abilities to the environmental filtering processes ^36^. However, the only similarity of phylogenetic pattern that persisted between species and family levels was that the phylogenetic measures of the Ninvnat community were found to be more influenced by the environmental variables, indicating the phylogenetic relatedness of the invasive and naturalized aliens across different taxonomic extents.

## Online Methods

### Data collection

We assembled China’s native and alien plant species from four nationwide checklists that provided the most up-to-date information on the Chinese flora. The species names were standardized, and infraspecific taxa and non-angiosperm species were removed. We further determined the alien species’ origin status (native or alien to China) and invasion status (introduced, naturalized, and invasive). The occurrence records (latitude and longitude) of these species were extracted from the Global Biodiversity Information Facility and the herbarium database of the National Specimen Information Infrastructure. All occurrence records were arranged by species names and cleaned for duplicates, points on the sea, coordinate uncertainties, and spatial autocorrelation. The cleaned occurrence records were arranged in a periodic interval of 20 years: until 1900, 1901-1920, 1921-1940, 1941-1960, 1961-1980, 1981-2000, and 2001-2021. At the end of these steps, we obtained a total of 8,65,752 occurrences for 19,674 species (native=19059, invasive=284, naturalized=197, introduced=134). We considered these species in this study (Fig.1a).

### Phylogeny construction and estimation of phylogenetic relatedness

We used a ‘mega-tree’ approach to generate a highly resolved phylogenetic tree by using the *V.PhyloMaker2* package ^37^ in R. We used the function build.nodes.1 to define the basal node of the genus as the most recent common ancestor of all the tips in the largest cluster of the genus and scenario 3 to add genera and species absent from the mega-tree ^38^.

In this study, we considered ecoregions as the community units having four categories of species (native, introduced, naturalized, and invasive). Using a custom script in R, we first identified the abundance of each species in 57 terrestrial ecoregions in China. The abundance values were used to calculate abundance-weighted standardized effect sizes (ses) as 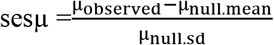, where μ is the phylogenetic measure. We used two phylogenetic measures in this study: mean phylogenetic distance (MPD) and mean nearest taxon distance (MNTD). The null distributions were built by randomly shuffling the species names across the tips of the phylogenetic tree 999 times. We used the *picante* package ^39^ in R to calculate the sesMPD and sesMNTD values. The sesMPD and sesMNTD values were further converted to the negative (−1 times) net relatedness index (NRI) and the nearest taxon index (NTI), respectively, the richness-standardized measures of MPD and MNTD. For NRI and NTI, positive values indicate phylogenetic clustering, whereas negative values indicate phylogenetic overdispersion ^35^.

The NRI and NTI values were estimated for each of the 57 ecoregions and seven periodic intervals. The ecoregions were further categorized based on possible community compositions having four categories of species (n=15 from 2^n^-1, where n = 4 species categories – native, introduced, naturalized, and invasive). Eight out of 15 community compositions were found to have NRI and NTI values (Fig.1b). The two phylogenetic measures (NRI and NTI) of these eight communities for seven periodic intervals were considered for further analysis.

### Data analysis

We compared the NRI and NTI values between the eight communities to test the dynamics of phylogenetic measures across the invasion continuum. We further characterized the phylogenetic changes with the introduction of alien species of three invasion categories as phylogenetic anomaly: 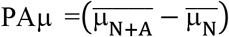, i.e., the difference between the phylogenetic measure (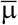; average for the ecoregions) between communities having native + alien species together (N + A) and native species alone (N). The PAμ values were compared with the null values and between the seven communities.

To test the first hypothesis (environmental filtering), we related patterns of phylogenetic relatedness to climate. The mean values of average annual temperature (BIO1) and annual precipitation (BIO12) for each of the ecoregions were considered as the explanatory variables in logistic regression models (by using the *glm* function in R) to check their influence on the variation in NRI and NTI values of the eight communities separately. To identify the effect of demographic variations on the phylogenetic relatedness, the ecoregions were categorized into four latitudinal (Lat1-Lat4) and seven longitudinal (Long1-Long7) grids at 10^0^N and 10^0^E intervals, respectively (Fig.1a). The phylogenetic measures across spatial scale (PS hereafter) at each of these grids were characterized as 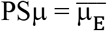, where 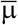 is the average phylogenetic measure of the ecoregions (E) within a grid. To test the second hypothesis (phylogenetic changes over time), we characterized Phylogenetic changes over time (PT, hereafter) for each community was characterized as 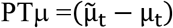, i.e., the deviation of the phylogenetic measure at each period (μ_t_, average for the ecoregions) from the median phylogenetic measure of the seven periods 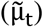. In addition to analyses at the species level, we conducted all analyses as outlined above for three families (taxonomic hierarchy) which had the maximum number of species, Asteraceae (n=1511), Fabaceae (n=972) and Poaceae (n=986), to test our third hypothesis (phylogenetic extent). Further, we compared the NRI and NTI values and the derived phylogenetic measures (PAμ and PSμ) between species and family levels. The methods and statistical analyses are described in detail in Supplementary Note 1.

## Acknowledgements

A.K.B. thanks the National Natural Science Foundation of China (grant 32050410299) and Guangdong Basic and Applied Basic Research Foundation (grant 2022A1515012015) for funding, and the Chang Hungta Science Foundation of Sun Yat-sen University for support. Y.H. thanks the National Natural Science Foundation of China (grants 41776166 and 42076117) and foreign cultural and educational experts project of the Ministry of science and technology (grant QNJ2021162001L) for funding. F.T. thanks the National Natural Science Foundation of China (grant 31200466) and the Natural Science Foundation of Guangdong Province, China (grant 2017A030313189). W.G. appreciates funding from the National Natural Science Foundation of China (grant 32160051). We thank Q. Li, Y. Jia, X. Lu, Y. Liu and H. Shao for help with data extraction.

## Data and Code Availability

The complete occurrence data set is provided as Supplementary Data 1. The R markdown files (having the R and Python scripts for data preparation and data analysis) and associated data are provided as Supplementary Software. Both Supplementary Data 1 and Supplementary Software are deposited in Figshare. All Supplementary Information are available on request.

---

**Extended Data Fig. 1:**
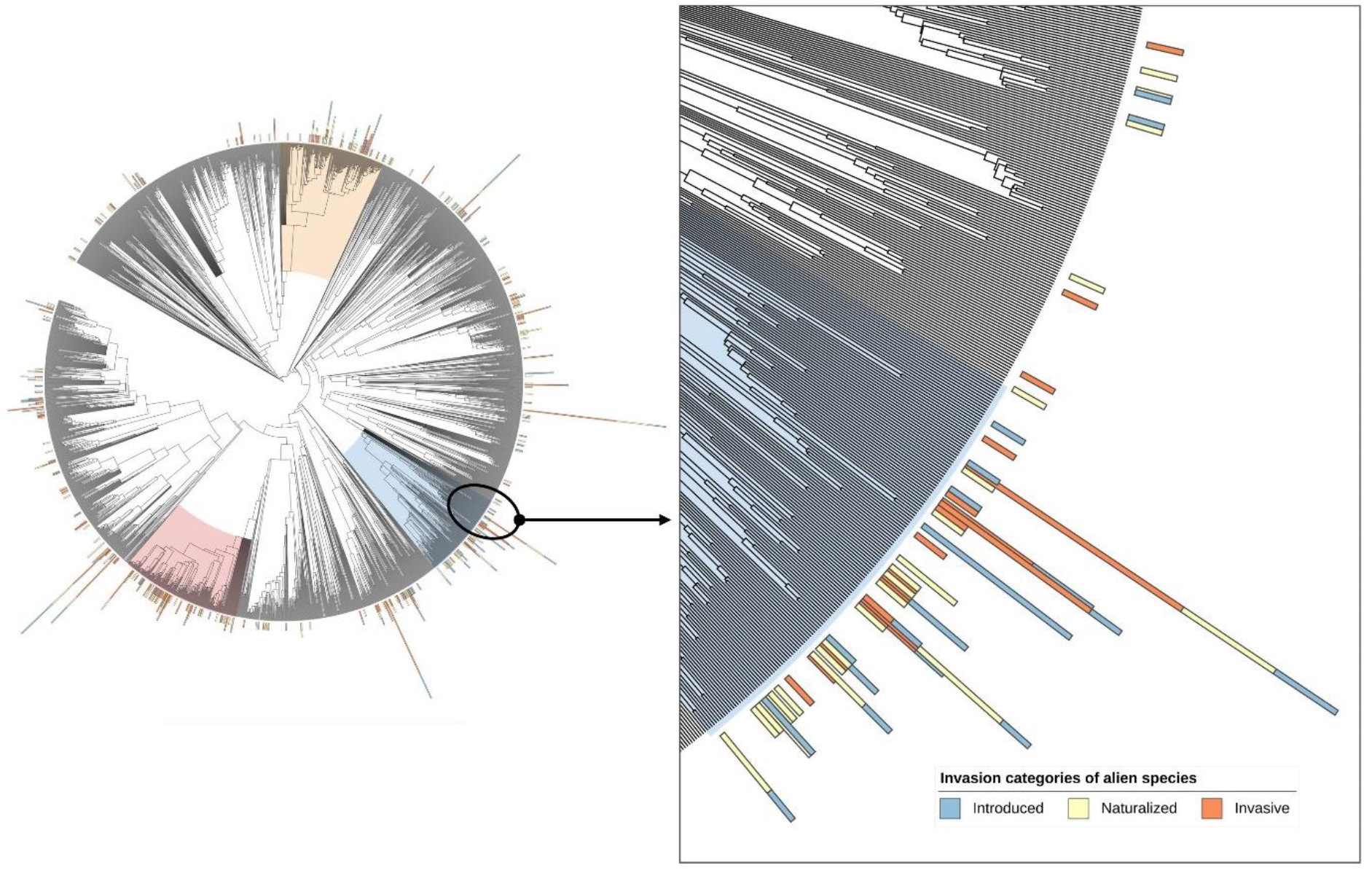
Genus-level phylogenetic tree showing phylogeny of the 2672 angiosperm genera. The stacked bars at the tip of the genus names indicate the number of species of three alien species categories. The clades of three major families analyzed separately in this study are shown.

## References

1 Simberloff, D. et al. Impacts of biological invasions: what’s what and the way forward. Trends Ecol. Evol. 28, 58–66 (2013).

2 Ricciardi, A. & Mottiar, M. Does Darwin’s naturalization hypothesis explain fish invasions? Biol. Invasions 8, 1403–1407, doi:10.1007/s10530-006-0005-6 (2006).

3 Daehler, C. C. Darwin’s naturalization hypothesis revisited. Am Nat 158, 324–330, doi:10.1086/321316 (2001).

4 Diez, J. M., Sullivan, J. J., Hulme, P. E., Edwards, G. & Duncan, R. P. Darwin’s naturalization conundrum: dissecting taxonomic patterns of species invasions. Ecol. Lett. 11, 674–681, doi: 10.1111/j.1461-0248.2008.01178.x (2008).

5 Thuiller, W. et al. Resolving Darwin’s naturalization conundrum: a quest for evidence. Diversity and Distributions 16, 461–475 (2010).

6 Qian, H. & Sandel, B. Phylogenetic relatedness of native and exotic plants along climate gradients in California, USA. Divers. Distrib. 23, 1323–1333, doi:10.1111/ddi.12620 (2017).

7 Blackburn, T. M. et al. A proposed unified framework for biological invasions. Trends Ecol. Evol. 26, 333–339, doi:10.1016/j.tree.2011.03.023 (2011).

8 Richardson, D. M. et al. Naturalization and invasion of alien plants: concepts and definitions. Divers. Distrib. 6, 93–107 (2000).

9 Omer, A. et al. The role of phylogenetic relatedness on alien plant success depends on the stage of invasion. Nature Plants 8, 906–914, doi:10.1038/s41477-022-01216-9 (2022).

10 Divisek, J. et al. Similarity of introduced plant species to native ones facilitates naturalization, but differences enhance invasion success. Nature Communications 9, 4631, doi:10.1038/s41467-018-06995-4 (2018).

11 Villalobos, F., Carotenuto, F., Raia, P. & Diniz-Filho, J. A. F. Phylogenetic fields through time: temporal dynamics of geographical co-occurrence and phylogenetic structure within species ranges. Philosophical Transactions of the Royal Society B: Biological Sciences 371, 20150220, doi:doi:10.1098/rstb.2015.0220 (2016).

12 Simberloff, D. & Von Holle, B. Positive interactions of nonindigenous species: invasional meltdown? Biological invasions 1, 21–32 (1999).

13 Donoghue, M. J. A phylogenetic perspective on the distribution of plant diversity. Proc Natl Acad Sci U S A 105, 11549–11555, doi:doi:10.1073/pnas.0801962105 (2008).

14 Gallagher, R. V., Randall, R. P. & Leishman, M. R. Trait differences between naturalized and invasive plant species independent of residence time and phylogeny. Conserv. Biol. 29, 360–369, doi:10.1111/cobi.12399 (2015).

15 Cadotte, M., Albert, C. H. & Walker, S. C. The ecology of differences: assessing community assembly with trait and evolutionary distances. Ecol. Lett. 16, 1234–1244, doi:https://doi.org/10.1111/ele.12161 (2013).

16 Fang, S. et al. Deterministic processes drive functional and phylogenetic temporal changes of woody species in temperate forests in Northeast China. Annals of Forest Science 76, 42, doi:10.1007/s13595-019-0830-2 (2019).

17 Axmacher, J. C. & Sang, W. Plant Invasions in China – Challenges and Chances. PLoS ONE 8, e64173, doi:10.1371/journal.pone.0064173 (2013).

18 Qian, H., Leprieur, F., Jin, Y., Wang, X. & Deng, T. Influence of phylogenetic scale on the relationships of taxonomic and phylogenetic turnovers with environment for angiosperms in China. Ecology and evolution 12, e8544–e8544, doi:10.1002/ece3.8544 (2022).

19 Qian, H. & Sandel, B. Darwin’s preadaptation hypothesis and the phylogenetic structure of native and alien regional plant assemblages across North America. Global Ecol. Biogeogr. 31, 531–545, doi:https://doi.org/10.1111/geb.13445 (2022).

20 Kuczynski, L. & Grenouillet, G. Community disassembly under global change: Evidence in favor of the stress-dominance hypothesis. Global Change Biol. 24, 4417–4427, doi:https://doi.org/10.1111/gcb.14320 (2018).

21 Wiens, J. J. & Donoghue, M. J. Historical biogeography, ecology and species richness. Trends Ecol. Evol. 19, 639–644 (2004).

22 Li, X.-H., Zhu, X.-X., Niu, Y. & Sun, H. Phylogenetic clustering and overdispersion for alpine plants along elevational gradient in the Hengduan Mountains Region, southwest China. Journal of Systematics and Evolution 52, 280–288, doi:https://doi.org/10.1111/jse.12027 (2014).

23 Pellissier, L. et al. Phylogenetic alpha and beta diversities of butterfly communities correlate with climate in the western Swiss Alps. Ecography 36, 541–550, doi:https://doi.org/10.1111/j.1600-0587.2012.07716.x (2013).

24 Crisp, M. D. & Cook, L. G. Phylogenetic niche conservatism: what are the underlying evolutionary and ecological causes? New Phytol. 196, 681–694 (2012).

25 Liu, H. et al. Strong phylogenetic signals and phylogenetic niche conservatism in ecophysiological traits across divergent lineages of Magnoliaceae. Scientific Reports 5, 12246, doi:10.1038/srep12246 (2015).

26 Cooper, N., Freckleton, R. P. & Jetz, W. Phylogenetic conservatism of environmental niches in mammals. Proceedings of the Royal Society B: Biological Sciences 278, 2384–2391, doi:10.1098/rspb.2010.2207 (2011).

27 Proches, S., Wilson, J. R. U., Richardson, D. M. & Rejmánek, M. Searching for phylogenetic pattern in biological invasions. Global Ecology and Biogeography 17, 5–10, doi:https://doi.org/10.1111/j.1466-8238.2007.00333.x (2008).

28 Lashchinskiy, N., Korolyuk, A., Makunina, N., Anenkhonov, O. & Liu, H. Longitudinal changes in species composition of forests and grasslands across the North Asian forest steppe zone. Folia Geobot 52, 175–197, doi:10.1007/s12224-016-9268-6 (2017).

29 Xie, Y., MacKinnon, J. & Li, D. Study on biogeographical divisions of China. Biodiversity & Conservation 13, 1391–1417, doi:10.1023/B:BIOC.0000019396.31168.ba (2004).

30 Hamrin, C. L. China and the challenge of the future: Changing political patterns. (Routledge, 2019).

31 Zhang, A., Hu, X., Yao, S., Yu, M. & Ying, Z. Alien, Naturalized and Invasive Plants in China. Plants 10, 2241, doi:10.3390/plants10112241 (2021).

32 Coutts, S. R., Helmstedt, K. J. & Bennett, J. R. Invasion lags: The stories we tell ourselves and our inability to infer process from pattern. Divers. Distrib. 24, 244–251, doi: 10.1111/ddi.12669 (2018).

33 Qian, H., Qian, S. & Sandel, B. Phylogenetic structure of alien and native species in regional plant assemblages across China: Testing niche conservatism hypothesis versus niche convergence hypothesis. Global Ecol. Biogeogr. 31, 1864–1876, doi:https://doi.org/10.1111/geb.13566 (2022).

34 Huang, W. et al. A well-supported nuclear phylogeny of Poaceae and implications for the evolution of C(4) photosynthesis. Mol Plant 15, 755–777, doi: 10.1016/j.molp.2022.01.015 (2022).

35 Webb, C. O., Ackerly, D. D., McPeek, M. A. & Donoghue, M. J. Phylogenies and Community Ecology. Annual Review of Ecology, Evolution, and Systematics 33, 475–505 (2002).

36 Weigelt, P. et al. Global patterns and drivers of phylogenetic structure in island floras. Scientific Reports 5, 12213, doi:10.1038/srep12213 (2015).

37 Jin, Y. & Qian, H. V. PhyloMaker2: An updated and enlarged R package that can generate very large phylogenies for vascular plants. Plant Diversity 44, 335–339, doi:https://doi.org/10.1016/j.pld.2022.05.005 (2022).

38 Qian, H. & Jin, Y. An updated megaphylogeny of plants, a tool for generating plant phylogenies and an analysis of phylogenetic community structure. Journal of Plant Ecology 9, 233–239, doi:10.1093/jpe/rtv047 (2015).

39 Kembel, S. W. et al. Picante: R tools for integrating phylogenies and ecology. Bioinformatics 26, 1463–1464, doi:10.1093/bioinformatics/btq166 (2010).

